# Bile Salts and Bacterial Bile Salt Hydrolase Activity Differentially Influence *Escherichia coli* Colonization and FXR Signaling in Colorectal Cancer Organoids

**DOI:** 10.1101/2025.09.20.676381

**Authors:** Ishita Dasgupta, Yixiao Ma, Durga Prasad Rangineni, Miriam Duran, Abhinav Bhushan

**Affiliations:** Department of Biomedical Engineering, Illinois Institute of Technology, Chicago, IL, USA

## Abstract

Bile salts are key regulators of host–microbe interactions and epithelial signaling in the intestine. Gut bacteria can deconjugate primary bile salts, which can be leveraged to engineer targeted therapies through bile salt hydrolase (BSH) activity. However, the interplay between bile salt signaling, microbial metabolism, and tumor-associated colonization remains poorly understood. Here, we establish an in vitro colon organoid platform to dissect bile salt–driven epithelial–microbe dynamics at cellular resolution. Using this system, we found that bile salts markedly modulated colonization of native *Escherichia coli* (EcAZ-2) and a BSH-expressing strain (EcAZ-2^LsBSH^) in healthy versus tumor murine organoids. Under bile salt exposure, EcAZ-2^LsBSH^ preferentially colonized tumor organoids, though overall colonization was lower than that of the non-BSH strain. Beyond effects on bacterial colonization, bile salts also directly activated host signaling pathways. In presence of bile salts, EcAZ-2LsBSH significantly enhanced activation of bile acid receptors such as farnesoid X receptor (FXR) by 2-fold in healthy organoids. These findings reveal a mechanistic bridge between in vivo microbial function and bile salt interactions, offering potential microbiota and bile acid targeted therapeutic strategies.

## Introduction

Colorectal cancer (CRC) remains a major global health concern, ranking as the third most lethal cancer in 2025. [1], [2] Imbalances in gut microbial communities are associated with inflammation, DNA damage, and immune dysfunction. These factors can act both as drivers and consequences of cancer development.[3,4] Further highlighting this connection, certain microbial communities such as *Fusobacterium nucleatum, Bacteroides fragilis, Escherichia coli (pks*^*+*^), can promote inflammation, induce DNA mutations and disrupt metabolic pathways thereby driving tumor growth [5],6].

Recent breakthroughs in three-dimensional cell culture technologies have led to the development of colorectal cancer organoids. These organoids offer a physiologically powerful and relevant model for dissecting the mechanisms underlying colorectal cancer progression [7]. Remarkably, these self-organizing organoids can proliferate without mesenchymal support[8]. Within organoids, stem cells expand and differentiate, develop forming structures that closely mimic the architectural feature of the intestine, including both crypt-like and villus-like regions populated with specific cell types [8,9]. Compared to two-dimensional monolayer cultures, organoids provide a more robust platform for modeling the physiology of gut lumen and investigating the crosstalk between the host and its gut microbiome [10]. Organoid technology holds immense potential, serving as a vital bridge the gap between cancer genetics and clinical trials. It complements conventional cell line and xenograft models, facilitating the development of personalized therapeutic strategies [11].

Numerous studies have leveraged colon organoids derived from tumors to explore cancer heterogeneity and evaluate drug sensitivity. A seminal study established a living biobank comprising patient-derived colorectal cancer organoids. This biobank not only preserves critical tumor characteristics but also enables personalized drug testing, marking a significant stride in precision medicine [11]. More recently, another compelling study reported the development of an optimized, growth factor-reduced culture medium that successfully maintains tumor-specific features in long-term colorectal cancer organoid cultures [7].

Despite these advancements, the crucial role of bile acids in organoid development remains largely underexplored. Bile acids are produced by the host and modified by the gut microbiome and regulate host metabolism and cellular signaling within the gut. Synthesized in the liver from cholesterol and conjugated for solubility, bile acids are secreted into the intestine where they engage closely with gut microbes [12]. Crucially, the interplay between bile acids and microbial colonization has not been sufficiently examined using organoid models [13]. Gut bacteria play a significant role in bile acid metabolism, primarily through BSH. BSH deconjugates primary bile acids, facilitating their conversion into secondary bile acids. These bile acids can then interact with the host receptors, such as FXR and Takeda G protein–coupled receptor 5 (TGR5), thereby influencing inflammation, epithelial proliferation, and ultimately, tumor progression [14].

Despite their profound impact on intestinal physiology and microbial ecology, bile salts are not routinely incorporated into organoid co-culture systems investigating host–microbe interactions. Notably, recent studies have reported conflicting effects of microbial bile acid metabolism on colorectal cancer: while one study found that bile acid mediated inhibition of FXR promotes cancer stem cell proliferation and tumor progression[15], another showed that bacterial BSH activity can exacerbate colorectal cancer through activation of immunosuppressive pathways and Wnt signaling [16]. These opposing findings highlight the need for physiologically relevant models to clarify how microbial bile acid transformations influence colorectal tumor biology. Gaining deeper insight into this relationship would enhance our comprehension of how bile salts and microbiota together influence the tumor environment and cancer progression.

To address this gap, this study hypothesized that: (1) bile salt exposure would differentially modulate the colonization capacity of native *E. coli* versus native BSH-expressing *E. coli* in both CRC and healthy colon organoids; (2) BSH activity would significantly alter host FXR and TGR5 signaling pathways; and (3) these interactions would reveal distinct gene expression signatures in healthy versus tumor-derived organoids. To investigate these hypotheses, we systematically examined the long-term colonization dynamics of gut bacteria in colorectal cancer organoids cultured in the presence of bile salts. We employed two bacterial strains previously characterized by Russell et al. [17] a bacterial chassis and carried no functional transgene, serving as an empty vector control (EcAZ-2) and an isogenic strain engineered to express BSH (EcAZ-2^LsBSH^). Crucially, EcAZ-2^LsBSH^ stably engrafts in the conventional mouse gut for the lifetime of the animal, retains BSH functionality, and systemically alters host bile acid profiles and metabolism [17]. By using this well-characterized tool in our organoid model, we directly dissected the specific epithelial responses to BSH-mediated deconjugation, providing cellular-level resolution that is challenging to achieve *in vivo*. [18]

## Materials and Methods

### Organoid culture

Healthy and tumor tissues from Apc^fl/WT^ CDX2^Cre^ mice were provided by Dr. Amir Zarrinpar’s Lab. Organoids were isolated from these tissues by enzymatic digestion using collagenase and gentamicin (1:1000 dilution), followed by washes with a wash media composed of Advanced DMEM/F12 (Gibco), 10% fetal bovine serum (Gibco), GlutaMAX (Gibco), and PenStrep (Gibco). Once the organoids were isolated, they were embedded in 60 µl droplets of Matrigel (Corning) and plated into pre-warmed 24-well plates. Matrigel was allowed to polymerize at 37 °C for 15-20 minutes before overlaying each with 500 µL of complete organoid culture media (see below).

Organoids were grown for five to seven days using the complete organoid culture media consisting of Advanced DMEM/F12, GlutaMAX, HEPES, and Penstrep, and supplemented with growth factors including B27-supplement (50X), N2 supplement (100X), Y-276322 (10 µM), Nicotinamide (10 mM), [Leu15]-Gastrin I (10 nM), N-acetylcysteine (1 mM), SB202190 (10 µM), A83-01(500 nM and recombinant mouse epidermal growth factor (EGF)(50 ng/ml) and L-WRN-conditioned media (50% volume/volume). Upon reaching confluency, the organoids were passaged in the ratio of 1:3.

### Bacterial culture

Two bacterial strains, EcAZ-2 and EcAZ-2^LsBSH^, were used in this study. EcAZ-2 is the native *Escherichia coli* strain isolated from the stool of a CR-WT C57BL/6 male mouse acquired from Jackson Laboratory (Bar Harbor, ME) and EcAZ-2^LsBSH^ is a genetically modified version of EcAZ-2 engineered to express the *Ligilactobacillus salivarius* (formerly *Lactobacillus salivarius*) bile salt hydrolase (BSH) enzyme [19],[17]. The addition of BSH to EcAZ-2 chassis did not alter its growth curve. Both strains were cultured in Luria Broth (LB) and incubated at 37 °C under standard aerobic conditions. Bacterial growth was assessed by measuring optical density at 600 nm (OD_600_) using a microplate reader (Biotek Synergy H1, Agilent).

To assess growth under bile salt exposure, overnight cultures of the both bacterial strains LB [19] medium supplemented with 100 µM each of tauro-β-muricholic acid (TβMCA; Cayman Chemical) and chenodeoxycholic acid (CDCA; Cayman Chemical). Cultures were loaded in triplicate on a 96-well plate. For the control we have only LB with bile salts, and it was also loaded in triplicate on the same plate. Optical density at 600 nm (OD_600_) was recorded every hour for 24 hours directly after 2 seconds of shaking using a microplate reader (Biotek Synergy H1, Agilent). The growth curve is provided in Supplementary Figure 1.

**Figure 1.**
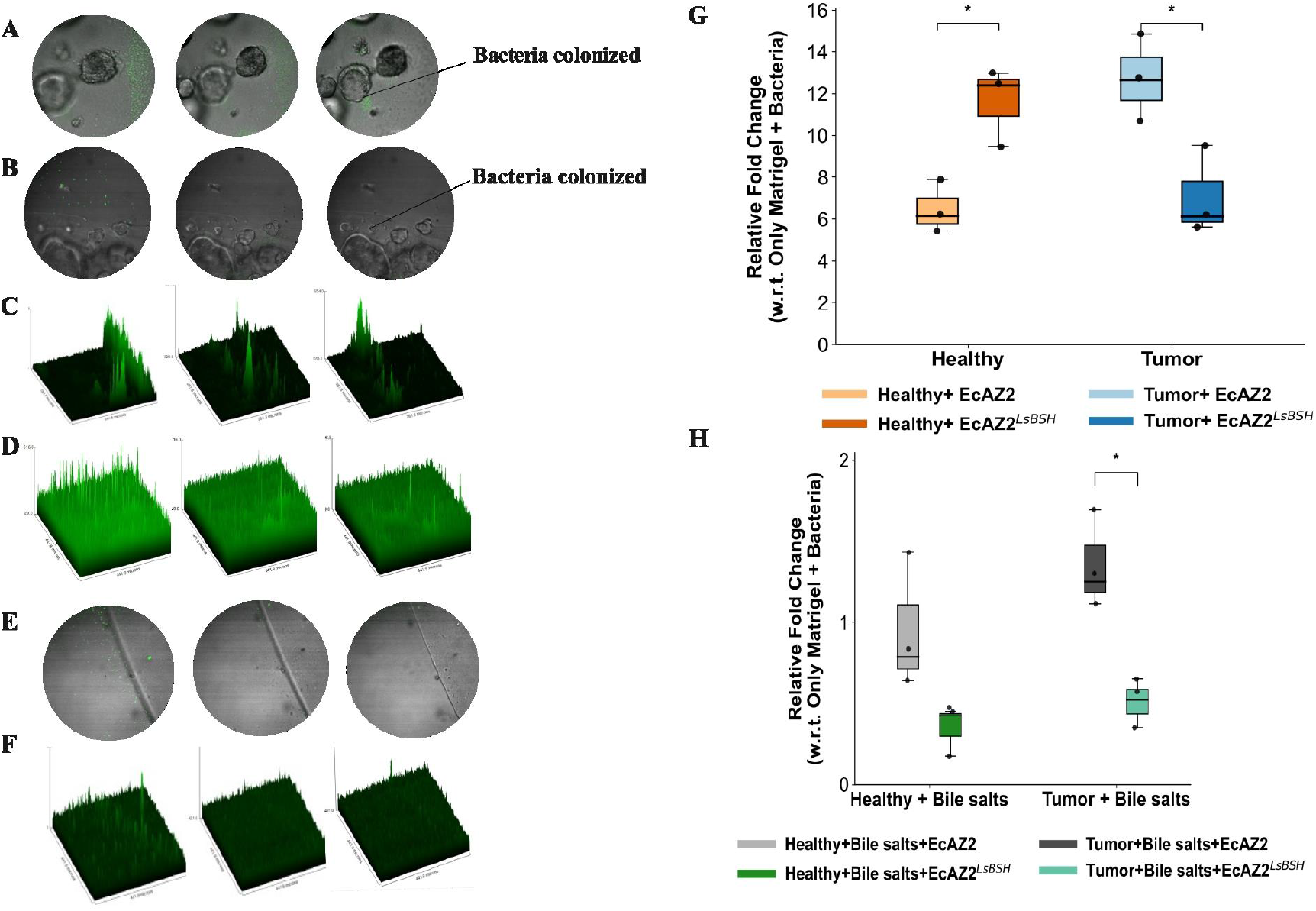
Gut bacteria colonization with the organoids **A.** Confocal microscopy z-stack images reflecting the EcAZ-2^LsBSH^ colonization in healthy organoids. GFP-expressing EcAZ-2^LsBSH^ are seen near the organoids in the first two images, with the third image showing the focal plane of EcAZ-2^LsBSH^ colonization with the organoids. **B.** Surface plots for the same z stacks images from A. The green, fluorescent peaks reflect the bacteria concentration localized at the plane of the z stack. **C.** Confocal microscopy z-stack images reflecting the EcAZ-2 colonization in healthy organoids. **D.**Surface plots for the same z stacks images from C. **E.** Confocal microscopy z-stack images reflecting the EcAZ-2^LsBSH^ colonization in control (only Matrigel) **F.** Surface plots for the same z stacks images from E. **G.** qPCR analysis on quantification of bacterial colonization with the healthy and tumor organoids. The relative fold change is measured with respect to colonization without bile salts. *p ≤ 0.05; **p ≤ 0.01; sample replicates (n=3). **H.**. qPCR analysis on quantification of bacterial colonization with the healthy and tumor organoids in presence of bile salts. The relative fold change is measured with respect to colonization without bile salts. *p ≤ 0.05; sample replicates (n=3).

### Colonization

Two experimental conditions were established to assess bacterial colonization: one in the absence of bile salts and the other in the presence of bile salts.

For the bile salt free condition, 24 hours after seeding the organoids, the media was replaced with complete organoid culture media without antibiotics. After 48 hr, 1 × 10^4^ colony-forming units (CFU) of bacteria were added to the culture media and incubated at 37^0^C with 5% CO_2_. The medium was replenished every 24 hours with fresh antibiotic-free complete organoid culture media, and incubation continued for up to 72 hours.

To investigate the effect of bile salts on colonization two bile salts namely tauro β muricholic acid (TβMCA, Cayman Chemical), and chenodeoxycholic acid (CDCA, Cayman Chemical) were used. The bile salts were prepared at a working concentration of 100µM in sterile deionized water. After 24 hr of organoids seeding, the media was replaced with antibiotic-free complete organoid culture media, supplemented with 100 µM CDCA and incubated for 24 hr. The bile salt concentration used in this study falls within the non-toxic range, as described in previous studies [17]. The media was then replaced with fresh antibiotic-free complete media containing 100 µM TβMCA and incubated for an additional 24 hr. Bacteria (1 × 10^4^ CFU) were subsequently added to each well. Colonization was visually assessed via microscopy for up to 72 hours; media was replaced daily. At the end of the incubation period, the organoids were lysed using RNA lysis buffer (Zymo) for mRNA extraction and qPCR analysis.

### Image analysis

The two bacterial strains were genetically modified via phage transduction to express green fluorescent protein (GFP) [17]. Imaging was performed using a confocal microscope (Nikon Eclipse Ti). Z-stack images and surface plots were acquired and analyzed using ImageJ (NIH).

### Quantitative Polymerase Chain Reaction (qPCR)

After 72 hr the organoids were washed with sterile phosphate-buffered saline (PBS) and lysed using the RNA lysis buffer (Zymo). Total mRNA was extracted using the Quick RNA Miniprep kit (Zymo) and mRNA concentration for each sample was quantified using Nanodrop (Thermo Fisher). Complementary DNA (cDNA) synthesis was performed using the qScript^TM^ cDNA Super Mix (Quanta Bio) on a Thermo Cycler C1000 Touch (BIO-RAD) following the manufacturer’s protocol. The primer sequences used are listed in the supplementary file Table 1.

For bacterial quantification, the organoids embedded in Matrigel were first dissociated using PBS-EDTA. Genomic DNA was extracted using the Monarch Genomic DNA Purification Kit (NEB), followed by qPCR analysis. The primers used for this analysis are listed in supplementary file Table 2.

### Relative gene expression analysis

Relative gene expression was calculated using 2^−ΔΔCt^ Method [20]. In brief, we first calculated the ΔCt of each target gene to the housekeeping gene -the ΔCt value is the difference between Ct of target genes and of housekeeping gene of each experimental and control group. By subtracting the ΔCt values of the experimental group from the control, ΔΔCt was calculated for each gene which was used to calculate the relative gene expression = 2^(−ΔΔCt)^. The relative fold change values were normalized by the average bacterial count per sample to adjust gene expression relative to bacterial load, providing an estimate of gene expression per bacterium. Normalized (bacteria-adjusted) expression values >1 indicate upregulation, whereas values <1 indicate downregulation.

### Data Analysis

For each group, organoids from three wells were analyzed. For the quantification of bacteria, a minimum of three independent samples were used for each group. Differences between experimental groups were compared using one-way ANOVA with Tukey’s post hoc test for multiple comparisons. Differences with a *p* < 0.05 were considered statistically significant.

## Results

### Assessing bacterial colonization dynamics in healthy and tumor organoids

Building on *in vivo* studies demonstrating the robust and persistent colonization of the murine gut by EcAZ strains [17], we first sought to characterize their colonization dynamics in our healthy and tumor-derived organoid models. Various *in vitro* methods have been employed to investigate the interaction between bacteria with intestinal organoids [21].These approaches include microinjection, organoid fragmentation, or introducing bacterial products directly to the media [21]. Since microinjection and fragmentation can cause structural damage to the organoids, for this study, we selected to introduce bacteria directly to the organoids embedded in Matrigel. Our approach controlled the bacterial load by adding a known number of bacteria. Regular media replacement helped prevent bacterial overgrowth and supported sustained colonization which was studied for up to 72 hours.

We quantified and compared bacterial colonization patterns between healthy and tumor organoids. Each organoid type was independently co-cultured with one of the two bacterial strains: native *E. coli* (EcAZ-2) and the genetically modified *E. coli* strain expressing BSH (EcAZ-2^LsBSH^). For each condition, a control consisting of only Matrigel with bacteria was included. Successful bacterial penetration of Matrigel and subsequent colonization of the organoids were consistently observed for both the strains **(Figure 1A & B)**. EcAZ-2 & EcAZ-2^LsBSH^ were clearly visible near the organoids, as confirmed by z-stack surface plots **(Figure 1C & D)**. We observed no colonization in the control condition, which consisted of Matrigel alone without the organoids **(Figure 1E & F)**. Relative colonization, measured against the control in healthy organoids for EcAZ-2 showed a 6-fold increase, while EcAZ-2^LsBSH^ exhibited approximately a 12-fold increase **(Figure 1G)**. In contrast, tumor organoids exhibited the reverse trend: EcAZ-2 colonization was upregulated by approximately 12-fold, whereas EcAZ-2^LsBSH^ showed only around 6-fold increase relative to the control **(Figure 1G)**. Our results showed distinct colonization patterns for the two strains depending on the organoid type, suggesting both strain-specific and host-specific interactions.

### Quantitative analysis of bacterial colonization in healthy and tumor organoids in the presence of bile salts

Subsequently, healthy and tumor organoids were pretreated with a combination of two bile salts at a physiologically relevant concentration of 100 µM prior to co-culture with bacterial strains. Fold changes in colonization were first calculated relative to Matrigel-only controls and then normalized to the corresponding condition without bile salts **(Figure 1H)**. Exposure to bile salts led to an overall reduction in colonization across both organoid types.

Notably, EcAZ-2^LsBSH^ exhibited higher colonization in tumor organoids compared to healthy ones when bile salts were present, though its colonization was still reduced relative to the non-BSH-expressing EcAZ-2 **(Figure 1H)**. While EcAZ2 preferentially colonizes the tumor organoids with bile salts (relative fold change >1) **(Figure 1H)**, EcAZ-2^LsBSH^ shows suppressed colonization. This suggests that the presence of BSH in EcAZ-2^LsBSH^ temporarily limits its growth and thereby reduces its colonization efficiency. These findings underscore the significance of bile salt exposure in influencing BSH activity, thereby shaping bacterial competitiveness and colonization potential.

While previous studies have explored how bile acids influence the composition and activity of the gut microbiome, the impact of bile acids on the colonization of gut bacteria remains largely unexplored [22]. Here, we examined how the bile salts CDCA and TβMCA influenced the bacterial colonization dynamics as well as BSH activity in EcAZ-2^LsBSH^. Given that CDCA acts as an FXR agonist and TβMCA as an antagonist, this approach allows us to investigate how bile acid deconjugation by the engineered bacteria impacts the colonization dynamics. The growth kinetics of both bacteria were assessed under bile salt exposure in absence of the organoids. The data indicates that both bile salts impaired bacterial growth irrespective of BSH activity (see Supplementary Figure 1).

### Influence of bacterial colonization on organoids in the absence of bile salts

To assess the impact of bacterial colonization on host transcriptional epithelial responses, we quantified the expression levels of key cell lineage markers: LGR5, CK20, and LYZ in both healthy and tumor organoids following colonization with the two bacterial strains. Fold changes were calculated within each condition, i.e., healthy organoids exposed to bacteria were compared with healthy organoids without bacteria, and tumor organoids exposed to bacteria were compared with tumor organoids without bacteria.

Expression values were normalized to bacterial load per sample to capture the per-bacterium transcriptional response, thereby accounting for inter-sample variability in colonization efficiency. LGR5 is an adult stem cell marker in intestinal crypts and is overexpressed in colorectal cancer [23]. CK20 is an epithelial differentiation marker commonly upregulated in colorectal cancer [24] with elevated expression observed in tissue and organoid samples from CRC patients [25]. LYZ is an antimicrobial peptide expressed by the Paneth cells with studies reporting its high expression during co-culture with bacteria [26]. Bacterial exposure, whether with commensal or pathogenic bacteria, can induce LYZ expression in intestinal cells [27].

Our findings show that in the tumor organoids, both bacterial strains significantly suppressed *Lgr5* expression (bacteria-adjusted expression < 1) relative to the no-bacteria control. However, after normalization, EcAZ2^LsBSH^ showed a significant increase in *Lgr5* expression compared to EcAZ2 **(Figure 2)**. *Ck20* expression was significantly elevated in tumor organoids colonized with EcAZ2^LsBSH^ compared to EcAZ2 **(Figure 2)**, suggesting that bacterial colonization with EcAZ2^LsBSH^ promotes enhanced epithelial cell differentiation within the tumor organoids.

**Figure 2.**
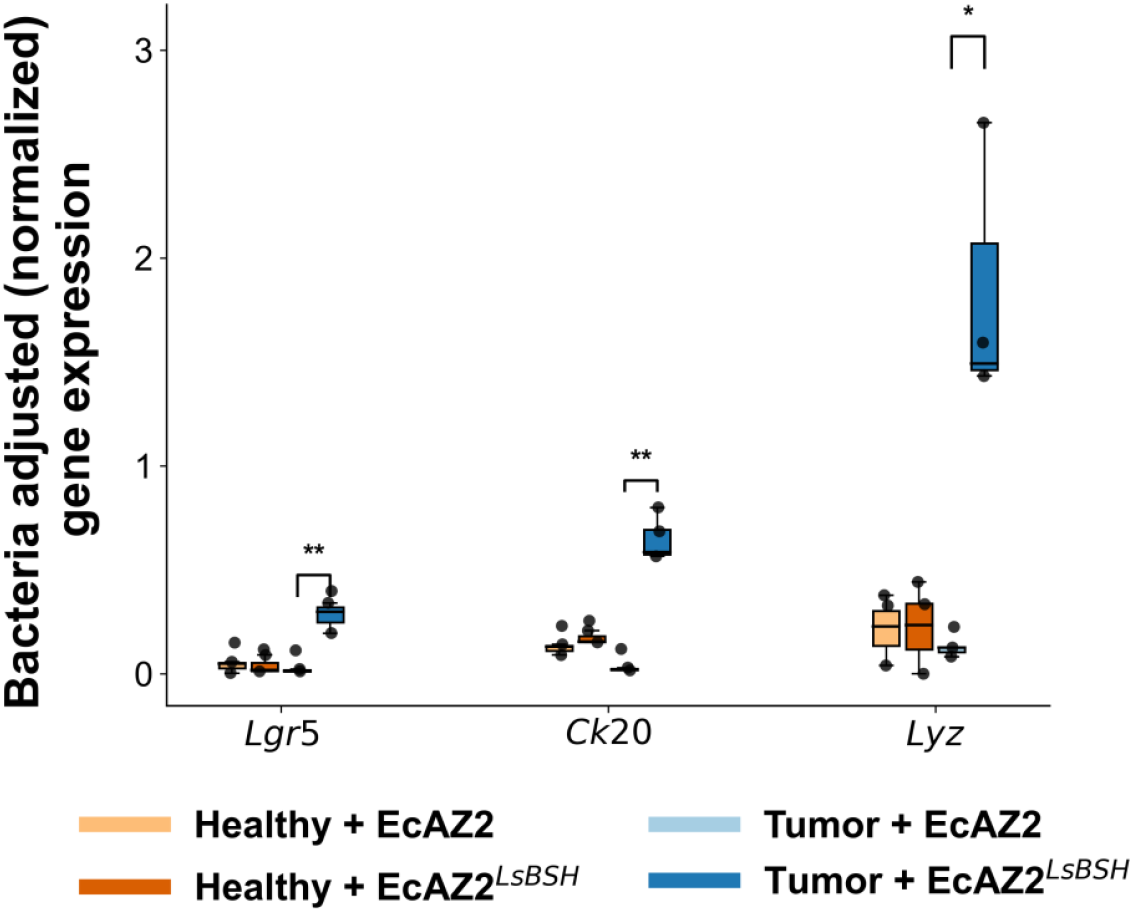
Normalized change in expression of *Lgr5, Ck20, and Lyz* in healthy and tumor organoids treated with *E. coli* strains EcAZ2 or EcAZ2^LsBSH^. Data are normalized to bacterial load per sample to account for colonization efficiency. EcAZ2^LsBSH^ significantly upregulated *Lgr5, Ck20 & Lyz* expression in tumor organoids (*, p < 0.05; **, p < 0.01).

*Lyz* expression was most strongly induced in tumor organoids treated with EcAZ2^LsBSH^ (bacteria adjusted gene expression ∼ 2) **(Figure 2)**. This pronounced normalized upregulation suggests a potential increase in Paneth cell activity or antimicrobial enzyme production triggered by engineered bacteria within the tumor organoids. These responses were not evident in the raw, unnormalized fold-change data (see Supplementary Figure 2), highlighting the value of normalization.

### Influence of colonization on organoids in presence of bile salts

Next, we examined the impact of bacterial colonization on host gene expression in healthy organoids cultured in presence of bile salts. Bacteria adjusted gene expressions were calculated relative to the control group (organoids cultured with bile salts and no bacteria) and normalized to the bacterial count colonizing the respective organoids in presence of bile salts **(Figure 3A)**.

**Figure 3A.**
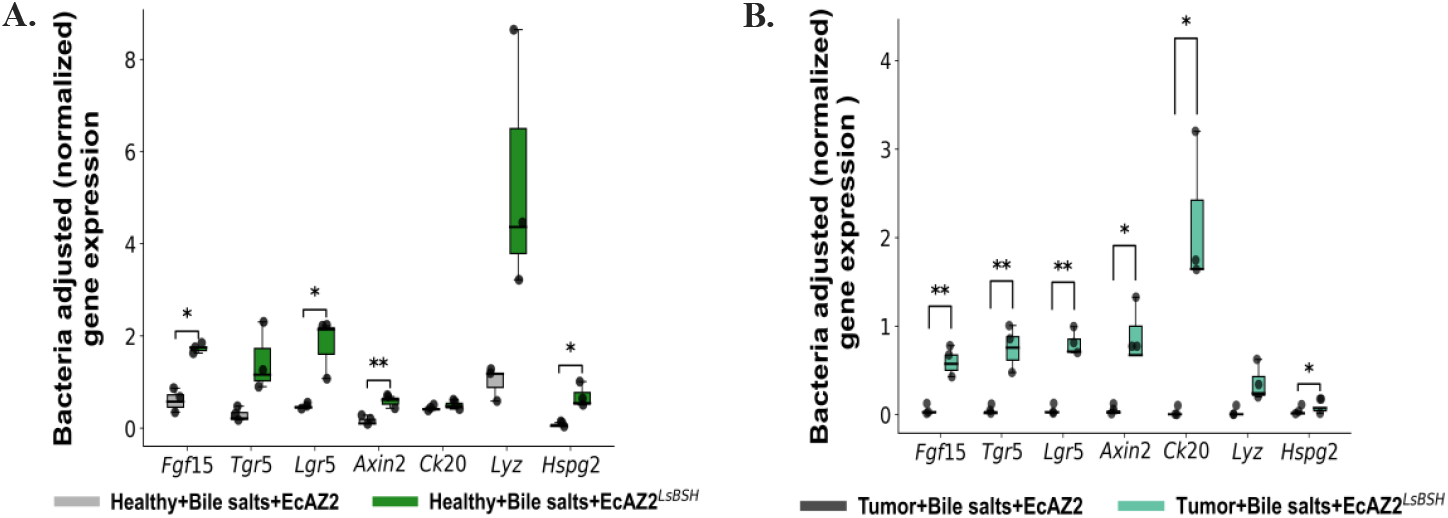
Bacteria-adjusted host gene expression in healthy organoids colonized with EcAZ2 and EcAZ-2^LsBSH^ following exposure to bile salts. **3B**: Bacteria-adjusted host gene expression in tumor organoids colonized with EcAZ2 and EcAZ-2^LsBSH^ following exposure to bile salts. The relative raw fold change is measured with respect to the control condition and normalized with the bacterial count colonizing the organoids in presence of bile salts. **p ≤ 0.01; *p<0.05

Our results demonstrated that EcAZ-2^LsBSH^ in presence of bile salts significantly upregulated the normalized gene expression of fibroblast growth factor 15 (*Fgf15)* by approximately 2-fold, implying enhanced bile acid deconjugation **(Figure 3A)**. In contrast, colonization with EcAZ-2 in the presence of bile salts did not change the expression of *Fgf15* **(Figure 3A)**. These results indicate that microbial deconjugation of bile by EcAZ-2^LsBSH^ significantly influences bile acid receptor pathways, thereby potentially influencing tumorigenesis in CRC organoids. In the colon, FXR activation induces the expression of the *Fgf15* gene, and its suppression has been linked to tumor development. [15.18]. Thus, FXR activity is inversely correlated with the development of CRC. To investigate the combined microbial and bile salts influence on FXR signaling, the organoids were treated with two bile salts prior to co-culture. In initial tests, we noticed a low-basal level of expression of *Fgf15* in the organoids. To test the suppression by EcAZ-2^LsBSH^, we decided to first upregulate the expression through an agonist. We activated FXR with CDCA, then inhibited it with TβMCA. This helped us to determine whether EcAZ-2^LsBSH^ would deconjugate TβMCA and reverse FXR inhibition and subsequently promote the expression of FXR-regulated genes, as well as those activated by other bile acid receptors, such as TGR5.

In a striking, bile-salt-dependent effect, colonization with BSH-expressing EcAZ-2^LsBSH^ triggered a nearly 2-fold increase in bacteria adjusted (normalized) expression of the stem cell marker *Lgr5* compared to the native EcAZ-2 strain **(Figure 3A)**. This effect was in stark contrast to the response observed in the absence of bile salts, highlighting a synergistic interaction between bile salt exposure and bacterial deconjugation in promoting a stemness of the organoids. Axis Inhibition 2 protein (*Axin2)*, another key marker associated with colon cancer stem cells and also well-established downstream target of the Wnt/β-catenin signaling pathway [28], showed a significant modest upregulation by EcAZ-2^LsBSH^ compared to EcAZ-2 **(Figure 3A)**.

Although both strains significantly suppress heparan sulfate proteoglycan gene 2 (*Hspg2*) expression overall, the EcAZ-2^LsBSH^ strain exhibits a slight increase in normalized expression relative to EcAZ-2. **(Figure 3A)**. *Hspg2*, which encodes the key extracellular matrix protein perlecan, is involved in cell adhesion and tissue integrity and is known to interact with histone-like protein A (*HlpA*) on the surface of *E. coli* [29]. Given that HLPA is a potential bacterial biosensor for host CRC signals,[30,31] monitoring *Hspg2* provides insight into host-microbe interactions that may facilitate bacterial attachment or persistence within the organoids.

The raw fold change qRT-PCR data without normalized has been provided in the supplement. (See Supplementary Figure 3). Normalization by the bacterial load mitigates the confounding effects of bile salts on bacterial growth and host responses. This helps us to understand that the adjusted gene expression changes are driven by bacterial activity from bile salt exposure alone; thus, ensuring that observed fold changes more accurately reflect per-bacterium effects on the host.

In tumor organoids, colonization by EcAZ-2 in the presence of bile salts led to a marked suppression in bacteria-adjusted (normalized) expression of most of the markers discussed above **(Figure 3B)**. In contrast, while EcAZ-2^LsBSH^ also downregulated these markers, its effect was less pronounced and, in some cases, paradoxical to its behavior in healthy organoids **(Figure 3A)**.

For instance, *Fgf15* was significantly downregulated by both bacterial strains **(Figure 3B)**, a stark reversal of the upregulation induced by EcAZ-2^LsBSH^ in healthy organoids **(Figure 3A)**. Concurrently, *Tgr5* was also downregulated, demonstrating that bile acid signaling and microbial bile acid modification in tumor organoids is markedly different from what we observed in healthy organoids.

Similarly, the stem cell markers *Lgr5* and *Axin2* were significant downregulation following colonization with either strain **(Figure 3B)**. While EcAZ-2^LsBSH^ induced a slight increase in expression of these markers compared to EcAZ-2, the bacteria-adjusted expression levels remained below baseline (bacterial adjusted expression<1).

Cytokeratin 20 *(Ck20)* was significantly upregulated, approximately two-fold, by EcAZ-2^LsBSH^, relative to EcAZ-2. Expression of *Hspg2* was also significantly reduced by both strains mirroring the pattern observed in the healthy organoids exposed to bile salts **(Figure 3A)**, with EcAZ-2 exerting a more substantial effect than EcAZ-2^LsBSH^. The raw (un-normalized) fold change qPCR data has been provided in the supplementary file. (See Supplementary Figure 4)

**Figure 4.**
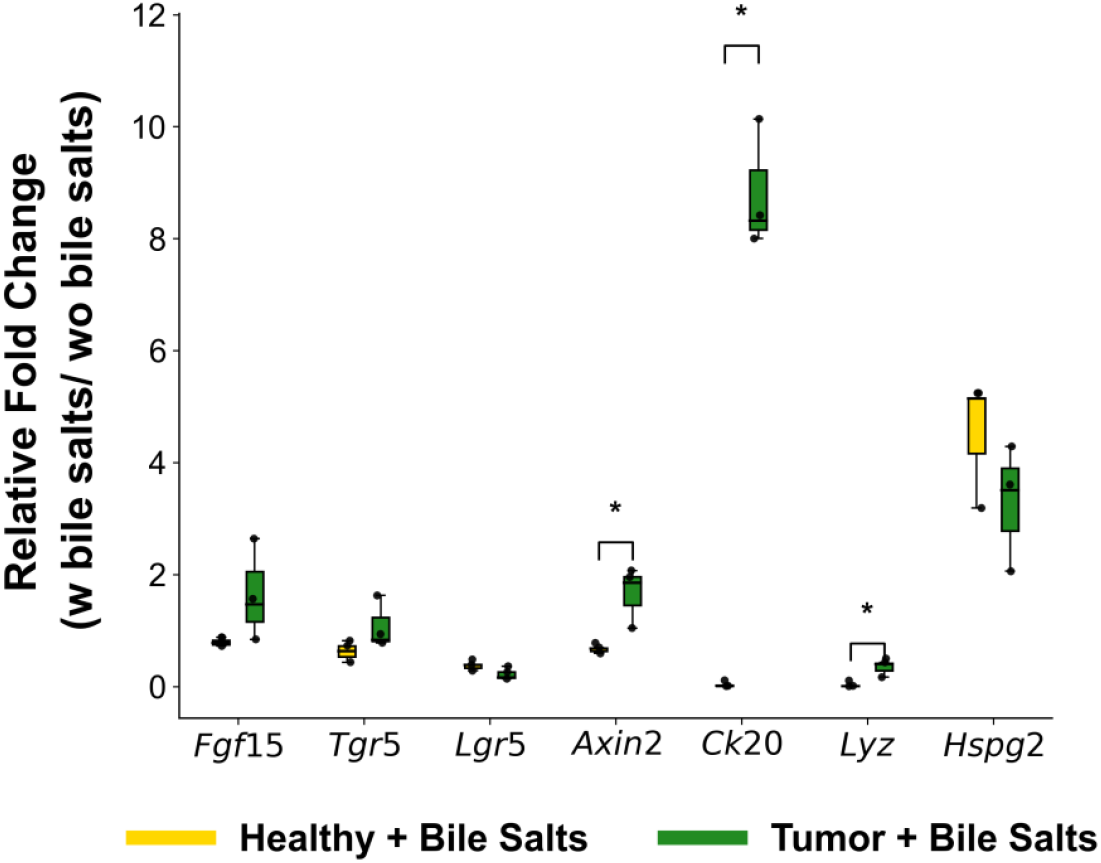
Relative fold change in gene expression in healthy and tumor organoids treated with bile salts. The relative fold change is measured with respect to the control condition comprising organoids not treated with bile salts. *p ≤ 0.05.

These findings suggest that EcAZ-2^LsBSH^ interacts less effectively with tumor organoids in a bile salt-rich environment apart from the effects on differentiation, or that unique regulatory factors within the tumor microenvironment override the expected host response. Further investigation will unravel the mechanisms underlying this context-dependent behavior.

### Baseline host gene expression in presence of bile salts without bacterial colonization

To deconvolute the effects of bile salts from those of bacterial colonization, we assessed host gene expression in organoids treated with bile salts alone, using untreated organoids as the baseline control. Here, the relative fold change is calculated relative to the organoids without any exposure to bile salts. The tumor organoids respond more strongly to bile salts than the healthy ones. Of all the markers tested, *Axin2* and *Ck20* have been significantly upregulated by approximately 2-fold and 8-fold, respectively **(Figure 4)**. This suggested exposure to bile salts activates the Wnt signaling pathway and differentiation of the stem cells in the tumor organoids. There was a significant increase in *Lyz* expression in the presence of bile salts. To establish a baseline for subsequent colonization experiments, we have also compared the constitutive gene expression profiles of tumor organoids to their healthy counterparts under bacteria-free conditions (See Supplementary Figure 5).

**Figure 5.**
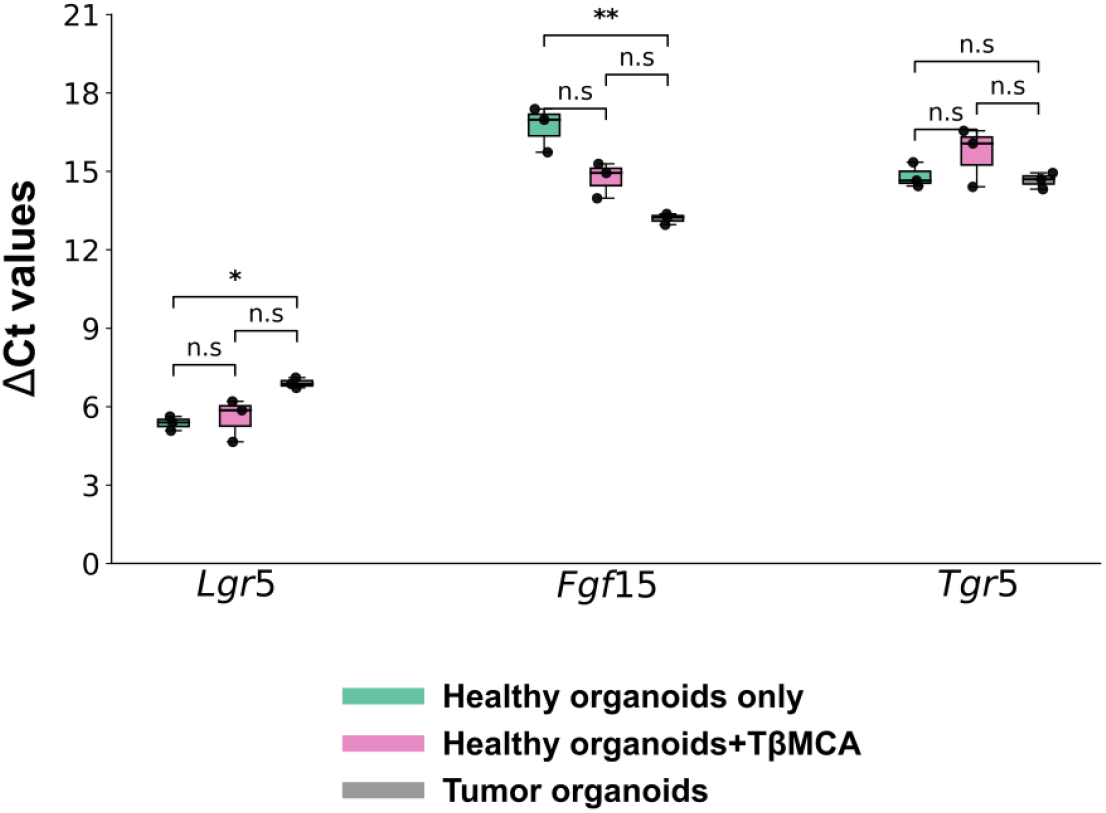
FXR inhibition in healthy organoids induces a tumor-like transcriptional profile. ΔCt values were calculated relative to the expression of a housekeeping gene (β-actin).

Upon treatment with bile salts, *Hspg2* expression was upregulated by approximately 6-fold in healthy organoids and 4-fold in tumor organoids **(Figure 4)**. However, these same genes were gets significantly suppressed in both organoids with bacterial colonization **(Figures 3A & B)**. This suggests that bile salts influence bacterial colonization dynamics by modulating host surface markers such as HSPG2. Overall, these findings demonstrate that bile salts exert profound and distinct effects on both healthy and tumor organoids, thereby acting as critical modulators of the host-microbe interface.

### Bile salt driven FXR inhibition aligns healthy organoids with tumor-like expression profiles

We then assessed whether FXR inhibition by bile salt would induce a tumor-like phenotype in healthy organoids. Upon TβMCA treatment, healthy organoids exhibited marked transcriptional changes consistent with a loss of FXR activity. Notably, expression of *Fgf15* was significantly downregulated, confirming effective inhibition of the FXR pathway. Concurrently, *Lgr5* was significantly increased. These changes, evident from ΔCt values, given that higher ΔCt values indicate lower gene expression, the observed increase in ΔCt confirms effective inhibition of FXR pathway.

Importantly, when we compared the ΔCt values of the markers (*Fgf15, Lgr5*, and *Tgr5*) between TβMCA treated healthy organoids and tumor-derived organoids, we found no statistically significant difference in their average ΔCt values **(Figure 5)**. Collectively, these observations reveal a direct mechanistic link between bile acid composition, FXR signaling suppression, and early molecular events associated with tumor initiation in the intestinal epithelium. This result supports the conclusion that FXR inhibition plays a vital role to reprogram healthy epithelial cells toward a tumor-like transcriptional state.

## Discussion

This study investigated the complex interplay between bile salts, bacterial colonization, and host epithelial responses using a physiologically relevant patient-derived organoid model. By integrating bile salts into the co-culture, we examined interactions between an engineered BSH-expressing *E. coli* strain (EcAZ-2LsBSH) and its native counterpart (EcAZ-2). This comparison revealed several key insights into the context-dependent nature of host–microbe interactions in both healthy and colorectal cancer organoids. Critically, our work builds upon foundational *in vivo* studies by Russell et al.[17], which established that this same EcAZ-2^LsBSH^ strain can stably engraft in mice and systemically alter host metabolism. Our organoid model provides a powerful, complementary platform to dissect the specific, cellular-level mechanisms driving these broader physiological phenomena. Our findings demonstrate that the microbial metabolism of bile acids, rather than bile acid exposure alone, is a primary driver of host signaling, and that the epithelial response to these interactions is fundamentally reprogrammed in tumor organoids compared to healthy ones. BSHs differ widely in substrate specificity and physiological effects depending on their bacterial origin. The BSH used in our study is distinct from other such enzymes, underscoring that other BSHs may elicit different host responses [32].

A central finding of our work is that BSH activity profoundly alters the intestinal microenvironment to the benefit of the colonizing microbe, but only in healthy organoids. In the presence of bile salts, BSH-expressing EcAZ-2^LsBSH^ demonstrated enhanced colonization and triggered a dramatic host transcriptional response, characterized by the upregulation of the stem cell marker *Lgr5* and the Paneth-like cell marker *Lyz*. This suggests a synergistic interaction where BSH-mediated deconjugation of bile salts creates secondary bile acids that, in turn, modulate host pathways to promote a stem-like state and enhance innate antimicrobial defenses. This complex phenotype potentially reflects active tissue maintenance and epithelial turnover. The activation of FXR signaling by EcAZ-2^LsBSH^ in this context further supports the hypothesis that microbially modified bile acids are potent signaling molecules that can orchestrate host physiology. These *in vitro* findings provide a direct cellular explanation for the systemic effects observed *in vivo*, where EcAZ-2^LsBSH^ was shown to alter circulating bile acid profiles and improve glucose tolerance [17].Our data suggests these systemic benefits may be initiated, at least in part, by the direct activation of protective FXR signaling at the epithelial level.

In stark contrast, the effects of BSH activity were blunted or even reversed in tumor-derived organoids. This finding is particularly significant when considering the challenges of translating live bacterial therapeutics into cancer therapy. While Russell et al.[17] demonstrated robust colonization of the healthy gut, other work has suggested that engineered probiotics may struggle to colonize the distinct niche of an established tumor *in vivo*. Our *in vitro* results, showing that tumor organoids are colonized but respond paradoxically, offer a potential mechanistic explanation for such discrepancies. The intrinsic dysregulation of signaling pathways in cancer cells, including the constitutive activation of Wnt signaling (as suggested by elevated baseline *Axin2* and *Lgr5*), may override or silence the signals initiated by microbial metabolites. This finding is critical, as it implies that therapeutic strategies aimed at modulating the microbiome-bile acid axis may have different, or perhaps limited, efficacy in established tumors compared to healthy tissue.

Our study also identifies *Hspg2* as a key, dynamically regulated factor at the host microbe interface. The significantly higher baseline expression of *Hspg2* in tumor organoids (See Supplementary Figure 5) could explain the enhanced colonization capacity of native *E. coli* in this environment, potentially by providing additional attachment sites. However, the subsequent downregulation of HSPG2 following both bacterial colonization and direct bile salt exposure suggests a sophisticated regulatory feedback loop. Bacteria may actively induce the shedding or suppression of this surface receptor to modulate host cell adhesion or evade host recognition. This complex interplay highlights HSPG2 as a potential therapeutic target for controlling bacterial association with tumors.

It is important to acknowledge the limitations of this study. As noted, discrepancies between our *in vitro* results and potential *in vivo* outcomes are themselves an important finding. While our model provides significant advantages, we did not directly quantify the profile of deconjugated bile acids in the culture medium. Future studies employing mass spectrometry would allow for a precise correlation between specific bile acid species, bacterial load, and the observed host response, strengthening the mechanistic links. Furthermore, our findings on EcAZ-2^LsBSH^ colonization *in vitro* differ from previous *in vivo* reports where it failed to colonize tumor tissue [17]. This crucial discrepancy underscores the complexity of the *in vivo* environment, where factors absent from our model such as a complex immune milieu, a structured mucus layer, and competition from a diverse commensal microbiota likely play a decisive role in determining colonization resistance. Exploring these differences is a vital future direction.

In conclusion, this work establishes a robust organoid-based platform to dissect the tripartite relationship between the host epithelium, microbiota, and bile acid metabolites. We demonstrate that bacterial BSH is a potent modulator of host stem cells and differentiation pathways in a manner that is fundamentally dependent on both bile salt availability and the health status of epithelium. The paradoxical responses observed in tumor organoids highlight the challenges and opportunities in targeting the microbiome for cancer therapy. Notably, treatment with bile salts particularly under conditions mimicking FXR inhibition was sufficient to shift healthy organoids toward a tumor-like transcriptional state, underscoring the central role of bile acid signaling in epithelial homeostasis. This model provides a valuable tool for future investigations into the mechanisms of bile acid and microbial-driven carcinogenesis and for the preclinical evaluation of next-generation engineered probiotics that have already shown promise in systemic metabolic disease models.

## Acknowledgements

We acknowledge receiving the bacterial strains and the tissue for organoid extraction from Dr. Amir Zarrrinpar and technical help from Dr. Arianna Brevi. This research was funded by National Science Foundation [2240045] and the National Institutes of Health [U01CA265719].

