## Supplemental file for "Bile Salts and Bacterial Bile Salt Hydrolase Activity Differentially Influence *Escherichia coli* Colonization and FXR Signaling in Colorectal Cancer Organoids"

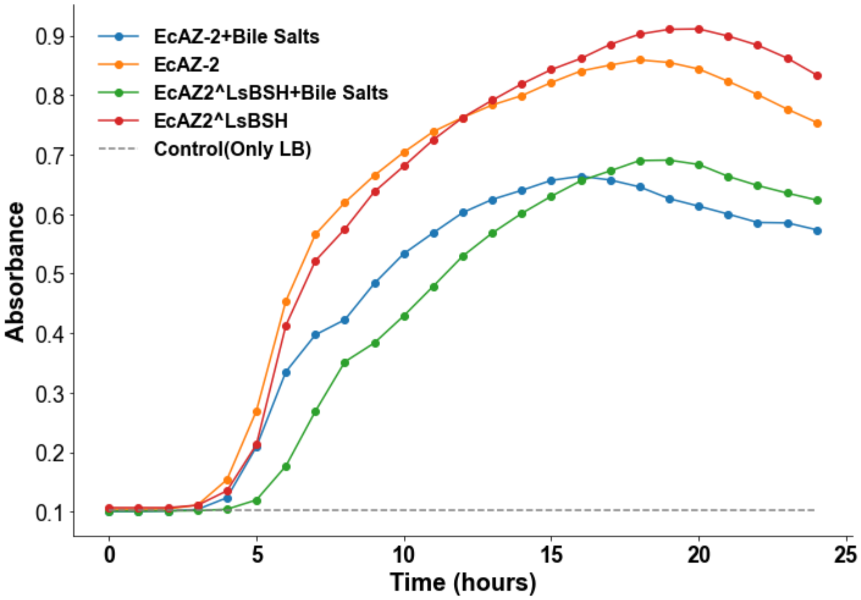


**Figure 1:** Growth curve of bacteria cultured in presence and absence of bile salts for 24 hours.

**Table 1:** qPCR primers for the mammalian markers tested on the organoids

| **Gene** | **Primer Direction** | **Sequence (5′→3′)** |
|---|---|---|
| β-actin | Forward | CCAGCCTTCCTTCTTGGGTAT |
| | Reverse | GGGTGTAAAACGCAGCTCAG |
| Lgr5 | Forward | AGAGCCTGATACCATCTGCAAAC |
| | Reverse | TGAAGGTCGTCCACACTGTTGC |
| CK20 | Forward | GGATTCGAGGTTCAAGTCACGG |
| | Reverse | TCTAGGTTGCGCTCCAGAGACT |
| LYZ | Forward | TGCCAGAACTCTGAAAAGGAATGG |
| | Reverse | CAGTGCTTTGGTCTCCACGGTT |
| FGF15 | Forward | ACGTCCTTGATGGCAATCG |
| | Reverse | GAGGACCAAAACGAACGAAATT |
| TGR5 | Forward | CACTGCTCTTCTTGCTGTGTTGG |
| | Reverse | GAGCGATAACAGAGTTCCAGGC |
| AXIN2 | Forward | ATGAGTAGCGCCGTGTTAGTGAC |
| | Reverse | CTTCGTACATGGGGAGCACTGT |
| HSPG2 | Forward | CATTCAGGTGGTCGTCCTCTCA |
| | Reverse | AGGTCAAGCGTCTGTCCTTCAG |
| SOX9 | Forward | CACACGTCAAGCGACCCATGAA |
| | Reverse | TCTTCTCGCTCTCGTTCAGCAG |
| RNF43 | Forward | CTGGCTATACCAGCATCGGACT |
| | Reverse | ATGCTGGCGAATGAGGTGGAGT |
| CDX2 | Forward | CATCAGGAGGAAAAGTGAGCTGG |
| | Reverse | TTTTCCTCTCCTTGGCTCTGCG |

The following *E.coli* bacterial primers were used for the quantification

| **Table 2:** qPCR primers for bacteria | | |
|---|---|---|
| *E.coli1* | Forward: | GCTACAATGGCGCATACAAA |
| | Reverse: | TTCATGGAGTCGAGTTGCAG |
| *E.coli* 2 | Forward: | AGAGCAAGCGGACCTCATAA |
| | Reverse: | TAGCGATTCCGACTTCATGG |

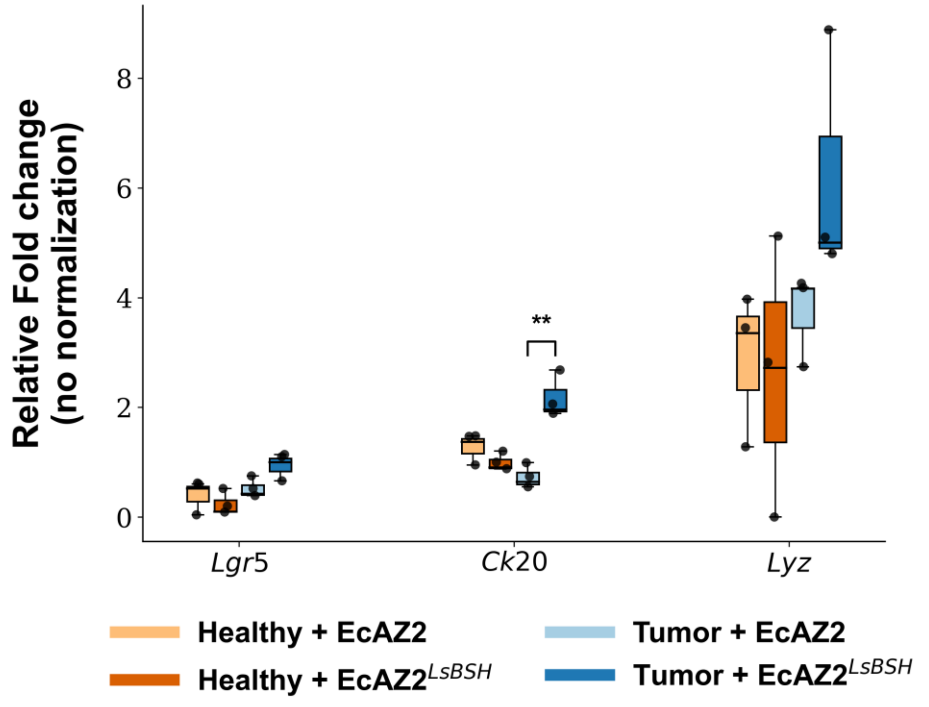


**Figure 2:** Fold change (w/o normalization) of organoids colonized in absence of bile salts**.**

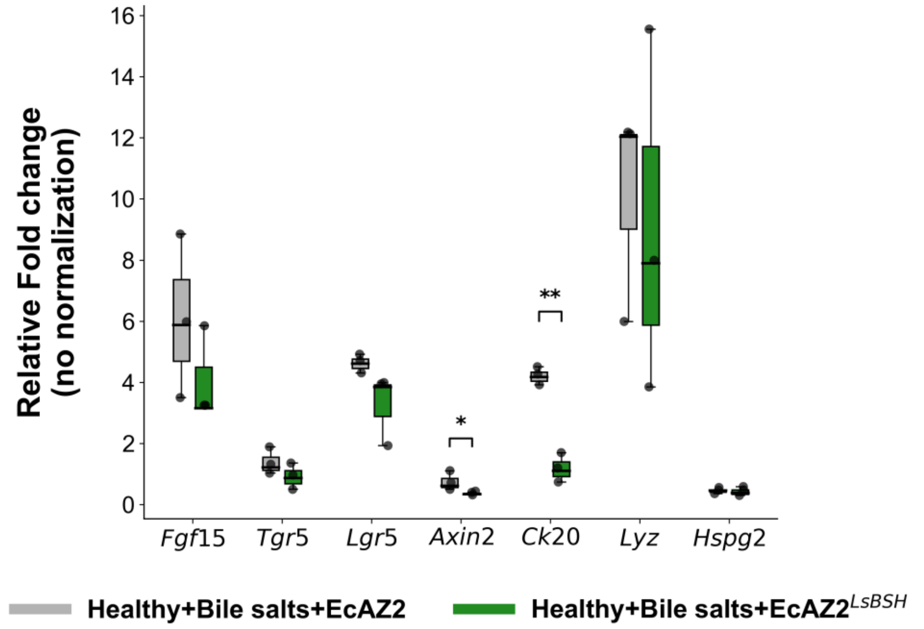


**Figure 3:** Fold change (w/o normalization) of healthy organoids colonized in presence of bile salts.

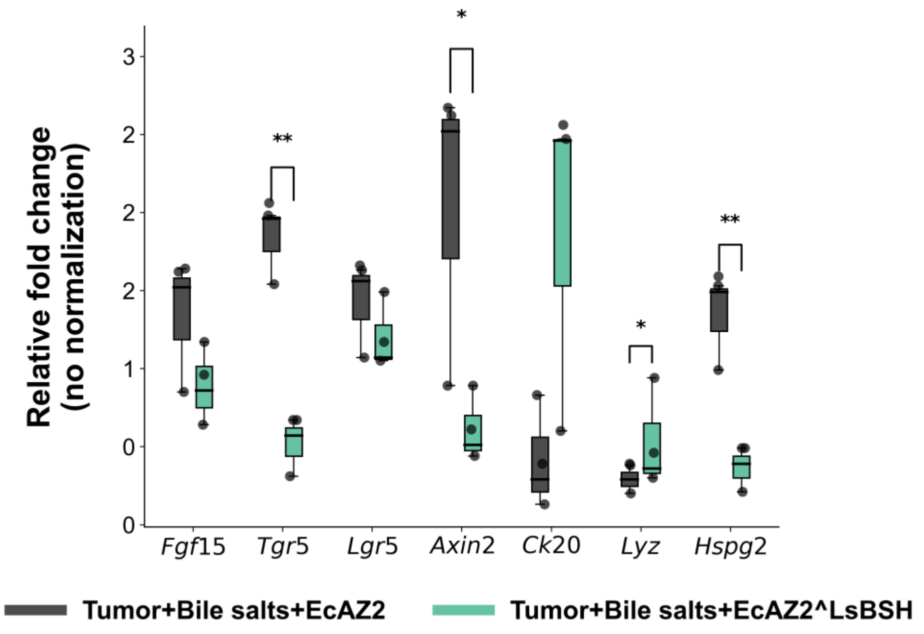


**Figure 4:** Fold change (w/o normalization) of tumor organoids colonized in presence of bile salts.

**Baseline host gene expression in the organoids without bile salts and bacterial colonization**

Using healthy organoids as the reference (fold change = 1), we identified significant intrinsic differences in the tumor organoid transcriptome that provide crucial context for interpreting responses to bacteria. We observed that the stem cell-associated genes Lgr5 and AXIN2 were constitutively upregulated in tumor organoids by 2-fold and 3-fold, respectively **(Figure 5)**. This pattern is consistent with previous studies on colorectal cancer tissues and organoids, where these genes are often overexpressed and associated with cancer stem cell properties. (van de Wetering et al., 2015). Interestingly, CK20, a marker of epithelial differentiation, showed significant marginal downregulation in tumor organoids **(Figure 5)**. Most strikingly, we observed a significant, approximately 10-fold upregulation of HSPG2 in tumor organoids compared to healthy ones **(Figure 5)**, which may explain the enhanced bacterial colonization observed in tumor organoids. This elevated baseline expression suggests that increased HSPG2 creates a more permissive surface for bacterial attachment. The Paneth cell marker LYZ, the proliferation marker SOX9, and the key CRC marker RNF43 were all significantly upregulated **(Figure 5)**. LYZ and RNF43 were significantly upregulated by approximately 2-fold and 3-fold in the tumor organoids **(Figure 5)**. These expression levels were consistent with previous findings reporting elevated levels of these markers in organoids derived from cancer tissues. (Tan et al., 2024; van de Wetering et al., 2015) It is well-established that RNF43 is frequently mutated and abundantly expressed in colorectal cancer tumor tissues compared to normal tissues. (Huang et al., 2023) Furthermore, SOX9 expression was increased by 4-fold in the tumor organoids **(Figure 5)**, a finding consistent with previous reports showing that its expression correlates with advanced stages of colorectal cancer and is more prominent in CRC tissues than in normal colon. (Zhou et al., 2020) Collectively, these observations suggest that our tumor organoids recapitulate established molecular traits of primary tumor tissue.

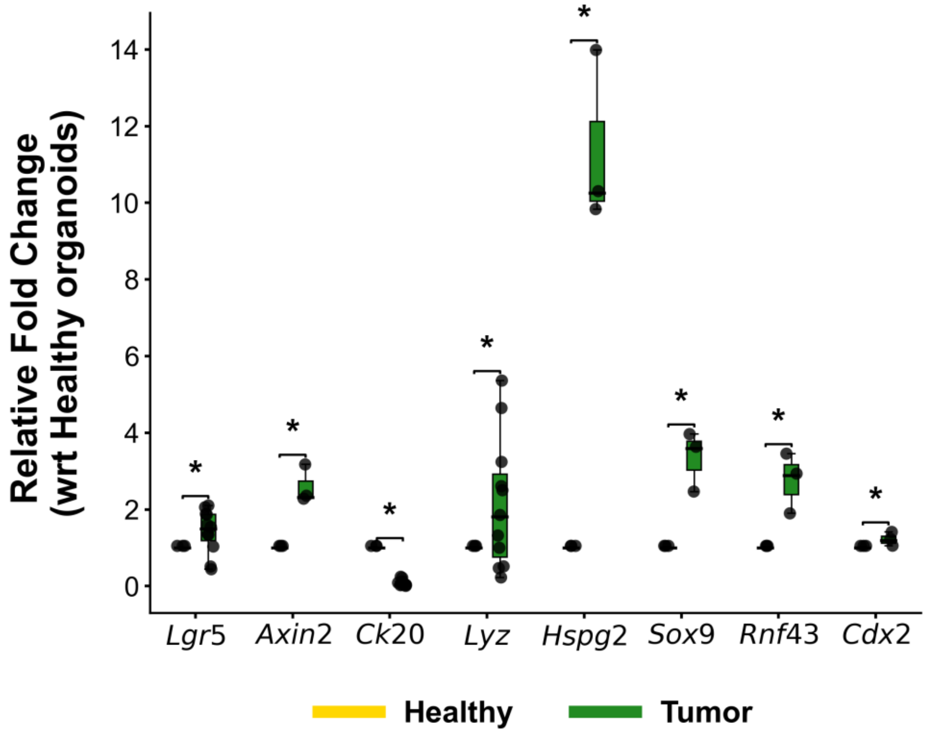


**Figure 5:** Gene expression data of healthy and tumor organoids in absence of bacterial colonization. The relative fold change for the tumor organoids is measured with respect to the healthy condition. $^{**}p \leq 0.01$; $^{*}p \leq 0.05$
